# Lipid vesicle formation by encapsulation of SMALPs in surfactant-stabilised droplets

**DOI:** 10.1101/2024.06.13.598947

**Authors:** Jorik Waeterschoot, Marta Barniol-Xicota, Steven Verhelst, Pieter Baatsen, Erin Koos, Jeroen Lammertyn, Xavier Casadevall i Solvas

## Abstract

Understanding the intricate functions of membrane proteins is pivotal in cell biology and drug discovery. The composition of the cell membrane is highly complex, with different types of membrane proteins and a huge variety of lipid species, Hence, studying cellular membranes in a complexity-reduced context is important to enhance our understanding of the roles of the different elements. However, reconstitution of membrane proteins in an environment that closely mimics the cell, like giant unilamellar vesicles (GUVs), remains challenging, often requiring detergents that compromise protein function. To address this challenge, we present a novel strategy to manufacture GUVs from styrene maleic acid lipid particles (SMALPs) that utilises surfactant-stabilised droplets as a template. Harnessing a new form of SMA linked to fluorescein, which we call FSMA, we demonstrate the assembly of SMALPs at the surfactant-stabilised droplet interface, resulting in the formation of GUVs when released upon addition of a demulsifying agent. The released vesicles appear similar to electroformed vesicles imaged with confocal light microscopy, but a fluorescein leakage assay and cryo-TEM imaging reveal their porous nature, potentially the result of residual interactions of SMA with the lipid bilayer. Our study represents a significant step towards opening new avenues for comprehensive protein research in a complexity-reduced, yet biologically relevant, setting.

## Introduction

Roughly one-third of all proteins in biological systems are categorised as membrane proteins.^1^ These proteins serve a wide array of functions such as transport, signalling, intracellular communication, and catalysis.^2,3^ Because of their roles in fundamental cellular processes, it is not surprising that membrane proteins are attractive drug targets. 60% of all FDA-approved targets are membrane proteins. ^4,5^ In a broader context, membrane proteins are characterised by their association with a lipid bilayer. Over the years, various experimental systems have been devised to facilitate their study. Notably, these include supported bilayers,^6–8^ proteoliposomes,^9^ lipid nanodiscs^10^ and giant unilamellar vesicles (GUVs).^11,12^ GUVs, in particular, hold special significance due to their cell-sized dimensions, with an inner aqueous phase separated from an outer phase. This unique setup allows researchers to investigate membrane proteins under more cell-like conditions, such as with distinct membrane potentials and concentration gradients.^13^ Furthermore, for the production of artificial cells with biologically-relevant properties, it is indispensable to develop effective strategies to accommodate the integration of a wide diversity of functional membrane proteins in GUVs.

Numerous protocols for the integration of membrane proteins entail the formation of small proteoliposomes with the aid of detergents.^14,15^ Various methods for generating GUVs from these proteoliposomes are available. One such method is the thin film hydration/electroformation technique, where proteoliposomes are deposited on a flat surface and subsequently rehydrated (with the application of an electric field in the case of electroformation).^16^ However, delicate proteins can be damaged by the drying process and detergents used. Additionally, there is a limited compatibility with the required buffer conditions for protein functionality.^11,17^ An alternative strategy involves fusing proteoliposomes with pre-formed GUVs. The amount of membrane protein that can be incorporated with this method is, however, limited.^13,18^ Another approach, developed by Weiss *et al.*,^19^ consist in encapsulating proteoliposomes in surfactant-stabilised droplets. The surfactant at the droplet interface induces vesicle fusion, leading to the formation of droplet-stabilised GUVs, which can later be released into an external buffer using a demulsifying agent like perfluoro-1-octanol (PFO). However, this method still necessitates the use of detergents in the initial proteoliposome preparation. Alternatively, an *in vitro* transcription/translation (IVTT) system can be encapsulated in a GUV to produce the membrane proteins and directly transfer them to the membrane. Nonetheless, this method requires the encapsulation of complex reaction mixtures, depends on the compatibility of all required reagents/protocols (for GUV formation and IVTT) and is not suitable for expressing all types of transmembrane proteins. ^20^

In an effort to circumvent the use of detergents during the isolation and purification of membrane proteins, researchers have devised innovative methodologies. For instance, Knowles *et al.*^21^ discovered the application of a styrene maleic acid (SMA) copolymer, which has the capacity to stabilise minute regions of lipid bilayers termed ‘lipid nanodiscs’ (or, in the case of styrene maleic acid: styrene maleic acid lipid particles, ‘SMALPs’). Notably, these techniques obviate the necessity for detergents, preserving the membrane proteins within their natural lipid milieu. The utility of SMALPs has been demonstrated across a broad spectrum of protein investigations,^22–25^ SMALPs can readily be generated from cell membranes or synthetic lipid membranes. The procedure entails the straightforward addition of SMA polymer to a vesicle solution, leading to the subsequent spontaneous insertion of its styrene groups into the lipid bilayer and self-assembly into a SMALP. This process results in the stabilisation of nanometre sized lipid patches, with the hydrophilic maleic acid groups oriented outward.^26^ However, it is important to note that while SMALPs offer numerous advantages, certain assays, such as investigation of the impact of transmembrane potential and concentration gradients, are not possible in these systems. Consequently, it would be of interest to transfer membrane proteins stabilised in SMALPs into GUVs, which offer well-defined intra- and extra-cellular-like environments.

In the present study, we explored the feasibility of leveraging the previously described droplet-stabilised GUV formation method^19^ to produce GUVs from SMALPs in lieu of traditional liposomes (see Figure 1 for an overview of the method). This would allow the transfer of membrane proteins to biomimetic GUVs while maintaining their original lipid environment, bypassing any harmful detergent mediated steps. To this end, we synthesised a novel fluorescent variant of SMA that we term FSMA. We first characterised this new polymer and utilised its fluorescent properties to further investigate the interaction between SMALPs and the droplet-stabilising surfactants that enable the conversion of lipid systems into GUVs. The released vesicles were then further analysed with (cryo-)TEM and a fluorescein leakage assay. Upon the realisation that the resulting GUVs were not effectively compartmentalising small molecules, several strategies to reduce GUV leakiness were further evaluated.

**Figure 1:**
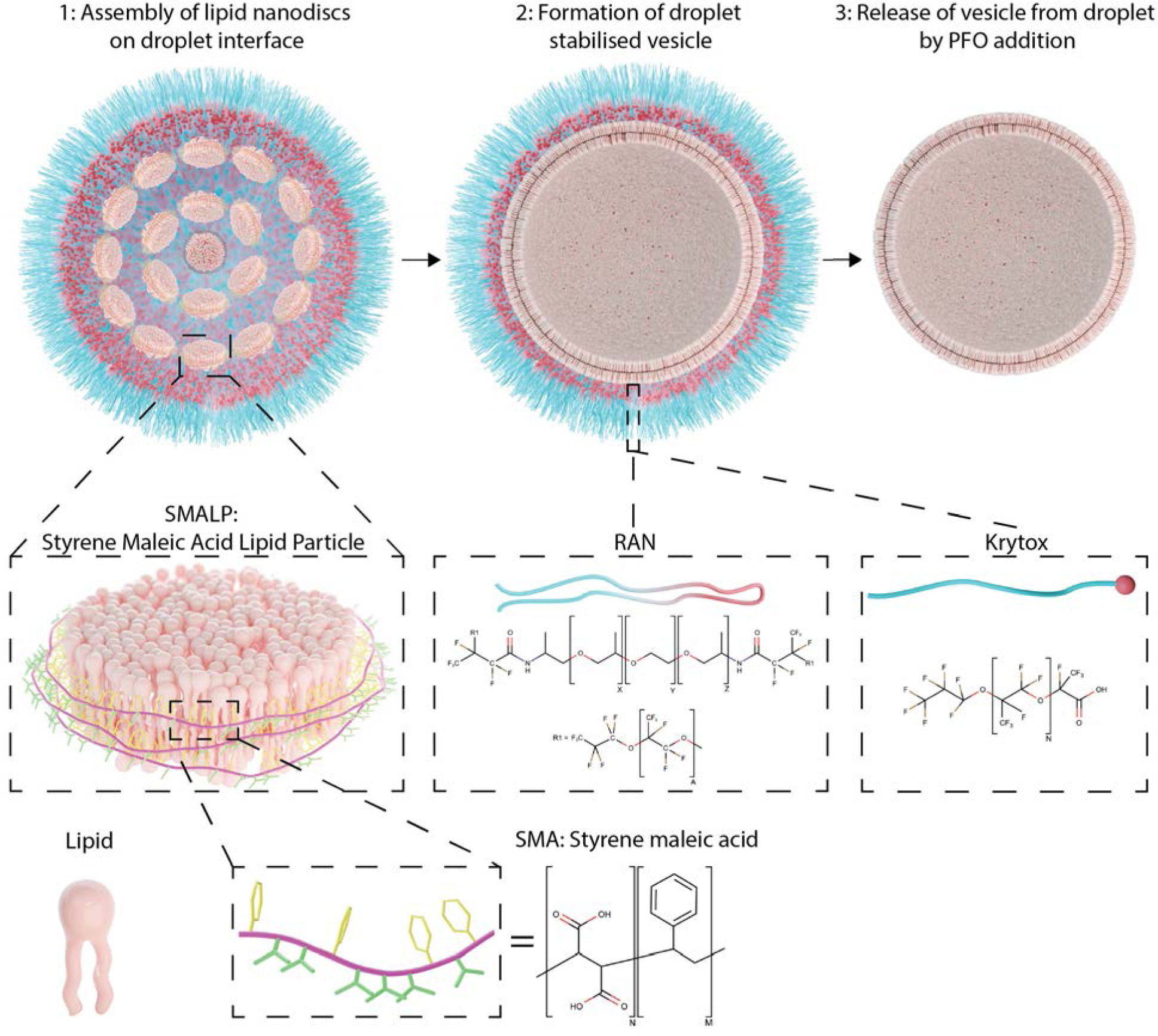
Schematic overview of vesicle formation from SMALPs.

## Results and discussion

### FSMA

For the formation of GUVs from SMALPs (via the surfactant-stabilised droplet method) the interactions between the SMALP components, mainly the SMA copolymer and the Krytox surfactant, needed to be investigated. Since standard SMA can not be tracked with optical microscopy, a new fluorescent variant (termed FSMA) was produced by linking SMA to a fluorescein fluorophore. FSMA was first characterised before the GUV formation was further investigated. For the chemical structure and the FSMA preparation, see Figure S1 and S2. This new polymer was first characterised via FTIR. As seen in Figure S3, the effective functionalisation of SMA with fluorescein was achieved, seen by the enlarged bands around 1600-1700 nm consistent with an increase of carbonyl groups.

The nanodisc-forming capability of FSMA was compared with that of SMA at different copolymer concentrations by solubilising neutral lipid vesicles (see Methods for the exact lipid composition). The total phosphorus content of the resulting FSMALPs/SMALPs was then quantified. As can be seen in Figure 2A, at lower FSMA concentrations (of up to 0.6 w/v%) the amount of lipids solubilised by FSMA is comparable to SMA. However, upon exceeding the threshold of 0.6 w/v% a reduction in the amount of solubilised lipids is observed, which can be suggestive of the aggregation of FSMA at higher concentrations, therefore reducing its lipid solubilisation potential.

**Figure 2:**
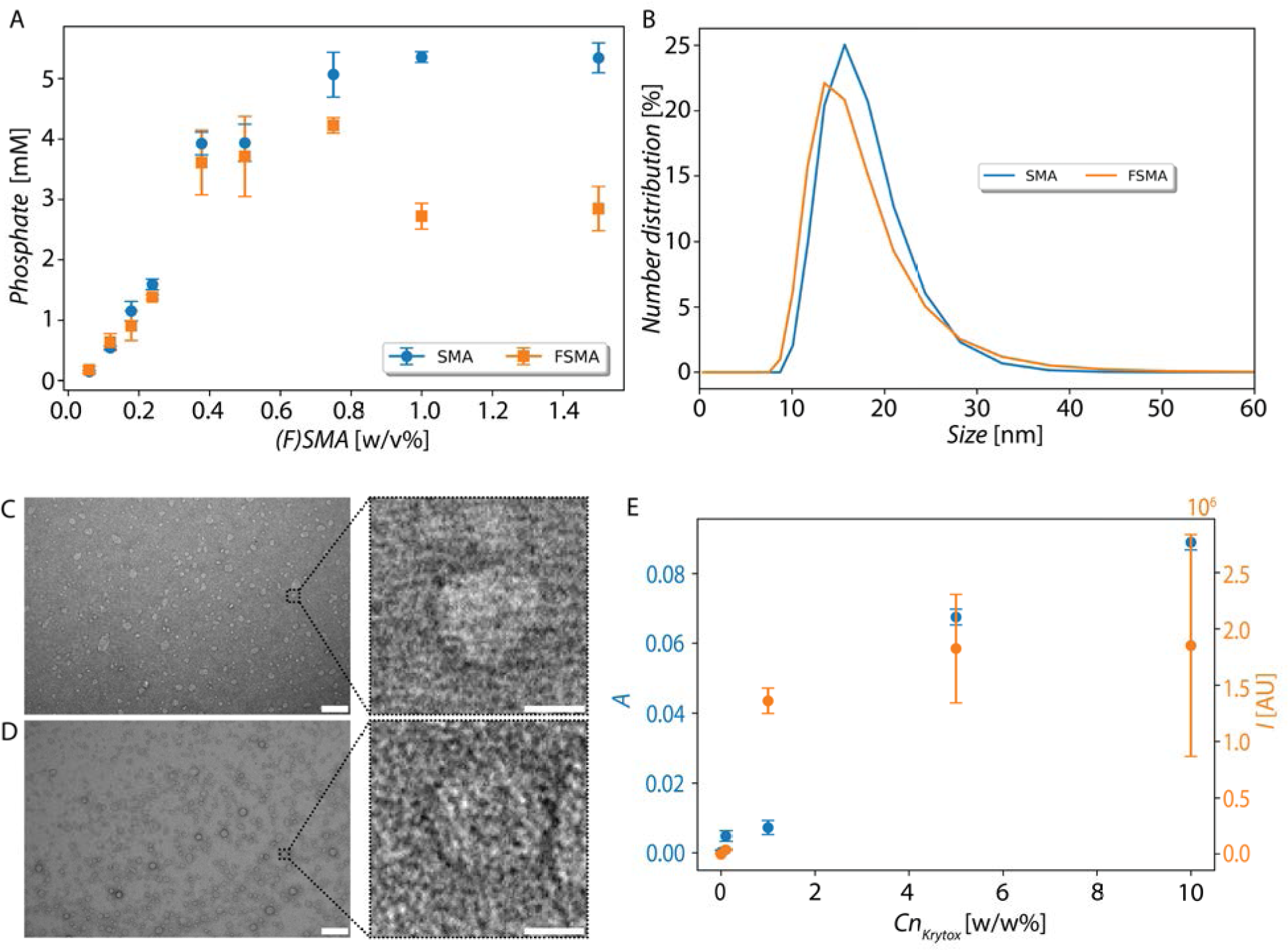
FSMA/SMA lipid solubilisation efficiencies, size analysis and interaction propensity with Krytox. A: FSMALP/SMALP formation efficiency from neutral lipid vesicle sample for SMA and FSMA. Experiments were performed in triplicate (technical and biological). Error bars represent one standard deviation. B: Number distribution of SMALPs and FSMALPs (neutral lipid composition, average of three). C: TEM image of SMALPs in B. D: TEM image of FSMALPs in B. C, D 100 nm scale bar, insets: 10 nm scale bar. E: Absorbance values (at 270 nm) measured in the oil phase on top of which an SMA polymer solution was incubated and fluorescence intensity values measured (470/550 nm, ex/em)in the oil phase on top of which an FSMA polymer solution was incubated (experiments performed in triplicate).

The sizes of FSMALPs/SMALPs (with neutral lipid composition) were analysed with DLS and TEM. Figure 2B-D confirms for both FSMALPs and SMALPs a size around 10-20 nm, similar to results found in literature.^22,27^

We next evaluated the capacity of FSMA to solubilise membrane proteins in comparison to SMA. To this end, red blood cells (RBCs) were subjected to FSMALP/SMALP formation. Band 3 was chosen given its prominence in the RBC membrane. ^28,29^ As a control for the specificity of membrane solubilisation, the resulting FSMALP/SMALP were also checked for the presence of, carbonic anhydrase 1, a common cytosolic protein. ^30,31^ FSMA and SMA both efficiently solubilised band 3 protein, as revealed by western blot analysis (Figure S4). In contrast, cytosolic carbonic anhydrase, being a soluble protein, was removed during the washing steps with low ionic and very low ionic strength (LIS and VLIS) buffers, and was not present in the FSMALP/SMALP samples (Figure S4). We conclude that the new FSMA behaves similar to SMA and, therefore, it can be used to provide more information on the mechanisms behind GUV formation from FSMALPs/SMALPs.

### Krytox and FSMA/SMA interaction

Vesicle formation in surfactant-stabilised droplets is mediated by the Krytox surfactant. As such, it was important to investigate whether there were any relevant interactions between this surfactant and the FSMA/SMA polymers at the droplet interface. For this, the polymer material (in SMALP buffer solution) was first incubated on top of an HFE 7500 oil phase containing varying amounts of Krytox surfactant. The absorbance (for the SMA) and the fluorescence (for the FSMA) were assessed in the oil phase after one day of incubation (see Figure 2E). Both results revealed an increase of the presence of the copolymer in the oil phase upon increasing Krytox concentrations, suggesting a direct interaction between the Krytox surfactant and the copolymer leading to its transfer into the oil phase. At 10 w/w% Krytox concentration, visible aggregation of the copolymer within the oil phase occurred, which could be attributed to the acidic nature of the Krytox molecules decreasing the pH of the copolymer solution.^32^ This, in turn, could explain the high variance of the results obtained in the experiments that used HFE oils with 10 w/w% Krytox, given the non-homogeneity of the resulting samples.

To further elucidate the potential interactions between different surfactant systems and fluorophores/copolymers, solutions of fluorescein isothiocyanate (FITC), FSMA and FSMALPs were encapsulated within droplets stabilised by these surfactants, illustrated in Figure 3. The FITC control shows a uniform distribution across all three different surfactant solutions, indicating that FITC does not engage in any significant interactions with Krytox molecules. In contrast, the results for FSMA show that the presence of Krytox induces FSMA accumulation at the droplet interface. In the last row of images in Figure 3, the accumulation of FSMALPs at the droplet interface is observed, which is a necessary step to form vesicles via the droplet-stabilised methodology.

**Figure 3:**
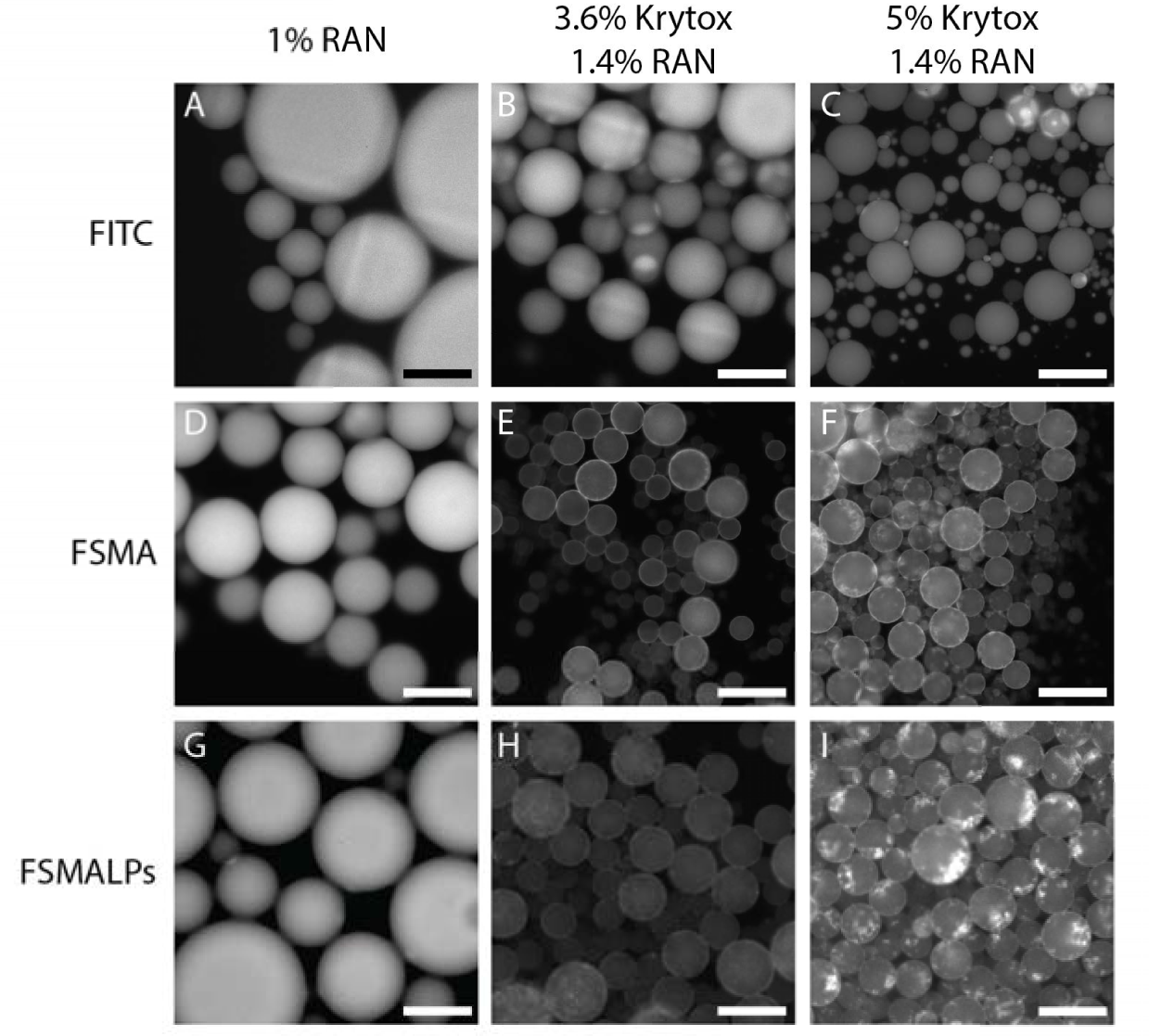
Fluorescence micrographs of droplets encapsulating FITC, FSMA and FSMALPs, stabilised by different surfactant mixtures. A, B and C: Droplets encapsulating FITC stabilised by 1 w/w% RAN, 3.6 w/w% Krytox/1.4 w/w% RAN and 5 w/w% Krytox/1.4 w/w% RAN respectively. D, E and F: Droplet encapsulating FSMA by 1 w/w% RAN, 3.6 w/w% Krytox/1.4 w/w% RAN and 5 w/w% Krytox/1.4 w/w% RAN respectively. G, H and I: Droplet encapsulating FSMA by 1 w/w% RAN, 3.6 w/w% Krytox/1.4 w/w% RAN and 5 w/w% Krytox/1.4 w/w% RAN respectively (100 µm scale bar).

The Krytox-FSMA interaction is potentially the result of a dimerisation of the carboxylic acid group present in the Krytox molecule and the maleic acid groups of the FSMA, ^33^ with the hydrogen bonds being potentially strengthened in the surrounding fluorinated medium.^34–36^ Additionally, the elevated concentration of sodium chloride serves to screen their charges, potentially enabling additional electrostatic interactions.

### Vesicle formation

To assess whether GUVs can be produced from SMALPs, we carried out a first set of vesicle release experiments. Here, droplets containing SMALPs made from four different lipid compositions (see Methods section) were produced using two different surfactant mixtures at two different concentrations. Representative images for the four different lipid compositions (before and after release from droplets) are shown in Figure 4A-H. As can be observed, SMALPs assembled at the droplet interface in all cases. It is important to highlight that, in some of the cases, aggregation of SMALPs into bright spots occurred. This was potentially the result of an acidifying effect of the Krytox molecules (containing carboxyl groups) leading to the destabilisation of the SMA copolymers. Importantly, though, GUVs were released for all four lipid compositions. Beside normally looking vesicles, a set of abnormal vesicles were observed, as shown in supplementary Figure S5. These were potentially the result of SMALPs that did not form GUVs, further interacting with the membrane of effectively formed GUVs, resulting in random formation of additional structures (such as lipid tether-like structures). To assess which were the most effective conditions for GUV production, the efficiency of GUV release from droplets was qualitatively assessed via fluorescence microscopy.

**Figure 4:**
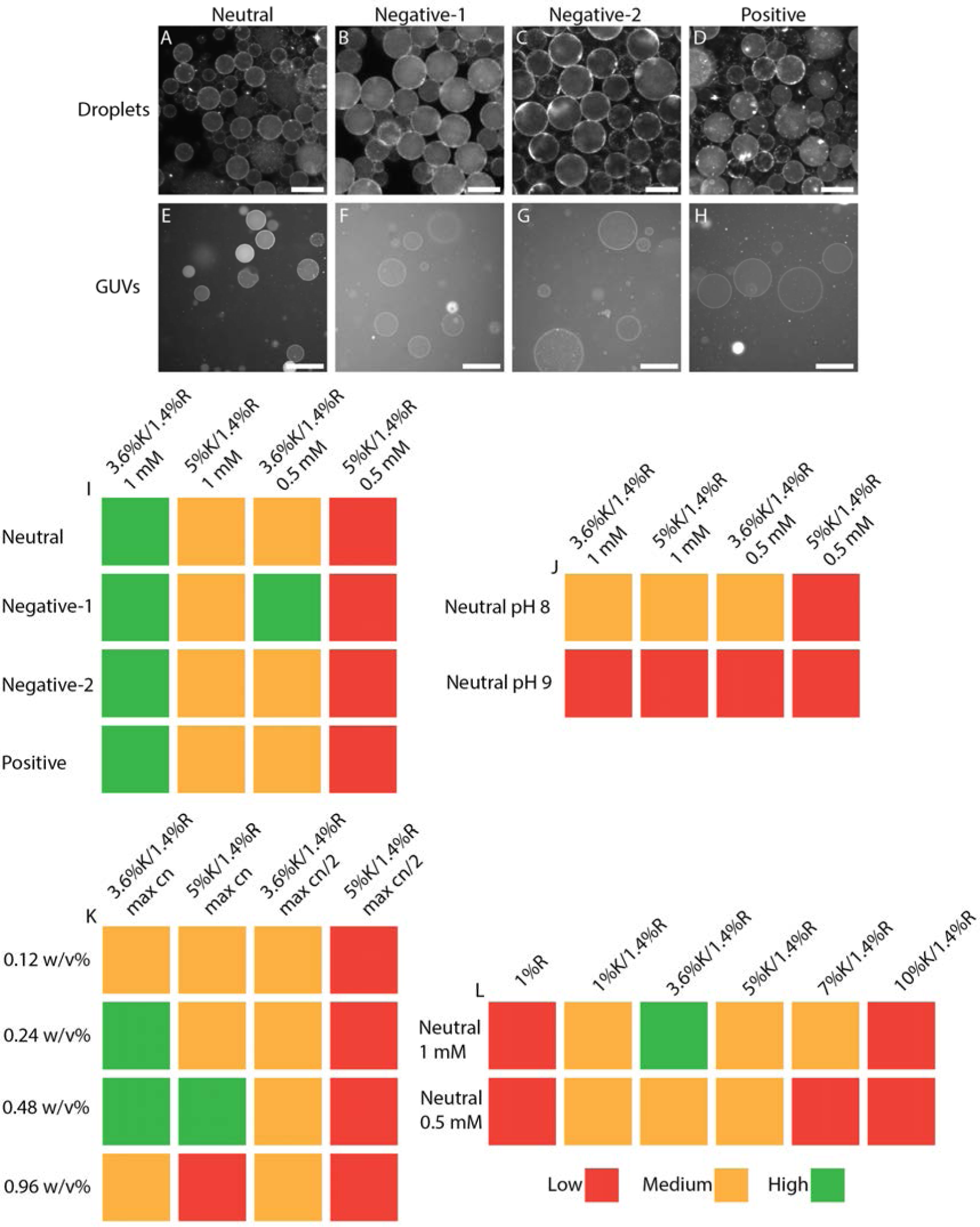
GUV release analysis of droplet stabilised GUVs formed from SMALPs. A-D: Droplets formed with 4 different SMALP solutions, 3.6 w/w% Krytox/1.4 w/w% RAN. E-H: Vesicles released from the droplets in A-D (100 µm scale bar). Qualitative release efficiencies in I-L were assessed by defining three different categories: low release (when just a few vesicles per imaging chamber were observed); medium release (when in every field of view a few vesicles were observed); and high release (when in every field of view many vesicles were observed). I: Qualitative release efficiencies for four different lipid compositions and two different lipid concentrations. J: Qualitative release efficiencies at pH 8 and pH 9. K: Qualitative release efficiencies for different SMA concentrations, max concentration indicates the use of the SMALPs directly after production without dilution, the max/2 samples were 1/2 diluted. L: Qualitative release efficiencies12for different Krytox concentrations.

As it is summarised in Figure 4I, the release efficiency was lower at the lower SMALP concentrations. In these cases, the droplets were not effectively stabilised by the surfactants and partial release of GUVs occurred spontaneously (without PFO addition). Therefore, in these cases the lipids contained in the SMALPs may not have had enough time to fuse together forming a fully enclosed, stable GUV. Additionally, it is also possible that the net lipid concentration to form a complete vesicle spanning the entire droplet interface was too low. Further investigations were performed towards the effect of pH, SMA concentration and Krytox concentration (see Figure 4J, K and L).

Lipid distribution in droplets with SMALPs at a higher pH are shown in Figure S9. At lower pH values (7.5) rings were observed, but this was accompanied by SMALP aggregation. At the higher pH of 9 the SMALPs were homogeneously distributed and did not form aggregated structures, but since no SMALPs assembled at the droplet surface, no ring formation was observed and no vesicles were released. The lack of vesicle formation in these conditions might be the result of two phenomena. On one hand, the carboxylic acid of the Krytox molecules at the droplet interface might not have been able to decrease the pH inside the droplet to an extent that the SMALPs were destabilised. On the other hand, the large deprotonation of the carboxylic groups may have resulted in a strong negative charge at the droplet interface, which was too large a charge to be screened by the sodium chloride present.

Neutral SMALPs formed with different amounts of SMA were encapsulated, imaged and afterwards released (see Figure S6 for images of the droplets and Figure 4K for the release efficiencies). SMALPs formed with low amount of SMA polymer resulted in unstable droplets. As discussed before, since droplets merged quickly the encapsulated SMALPs only had a limited time to interact and formed a limited amount of vesicles, explaining the average release efficiencies. At higher SMA concentrations, droplets were more stable and rings were observed, indicating possible vesicle formation and explaining the higher GUV release efficiencies in these conditions. At the highest polymer concentration strong aggregation was observed, typically into one fluorescent clump. Here, almost no vesicles were released. The high concentration of free SMA potentially destabilises the vesicles during the release process by breaking up the vesicles in smaller lipid structures.

The effect of the Krytox concentration in the oil phase was investigated for neutral SMALPs at a 1 mM and at 0.5 mM lipid concentration. Fluorescence micrographs of the droplet formation are shown in Figure S7 and S8, and the release efficiencies are displayed in Figure 4L. In case of the pure RAN surfactant, homogeneous distribution of SMALPs in the droplets were observed, indicating no interaction between RAN and the lipid SMALPs and, consequently, no formation and release of GUVs. Release of vesicles was possible at intermediate Krytox concentrations and highest at around 3.6 w/w% Krytox/1.4 w/w% RAN. At lower concentrations release was poorer, possibly as a result of insufficient assembly of SMALPs at the droplet interface and no vesicles being formed. At the higher Krytox concentrations, the acidifying effect of the Krytox was so large that it potentially led to an excess aggregation of SMALPs, thus disturbing the vesicle formation process. Additionally, at these high concentrations the interaction between the Krytox and the SMA in the vesicles might have become strong enough to tear apart the vesicles during the release process. ^37^

In order to ascertain the formation of a complete lipid bilayer, as opposed to the simple aggregation of lipid SMALPs, we employed negative stained transmission electron microscopy (TEM) to examine the final released vesicles. The TEM images revealed the presence of vesicle-like structures in which individual SMALPs are no longer distinguishable. Furthermore, these vesicles closely resemble electroformed vesicles, which provides compelling evidence of the fusion of lipid SMALPs into a more homogeneous lipid bilayer system (see Figure 5A and C).

**Figure 5:**
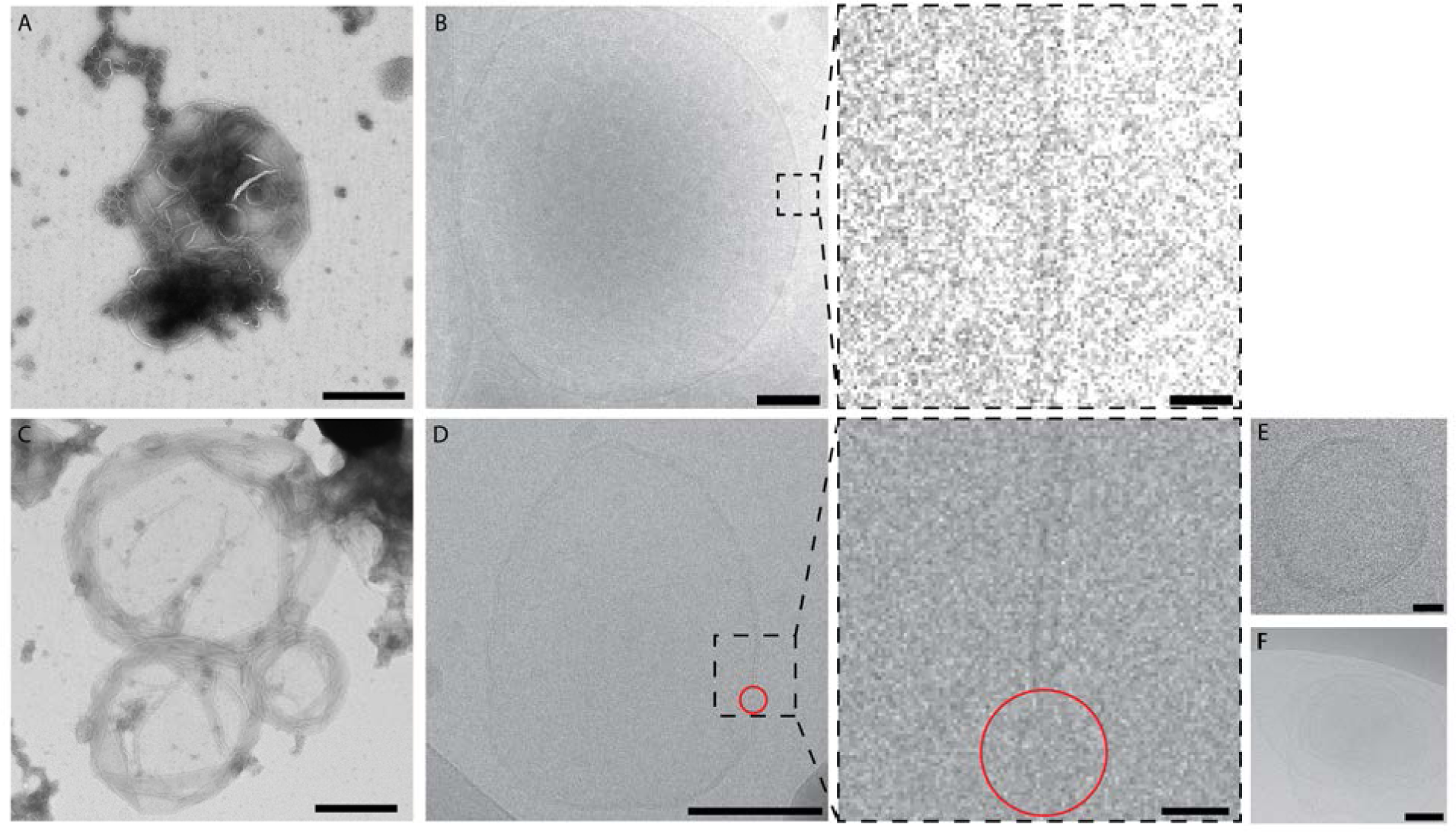
Electron micrograph images of both electroformed and SMALP formed GUVs. A: TEM micrograph of electroformed vesicles (500 nm scale bar). B: Cryo-TEM micrograph of vesicle formed via electroformation (100 nm scale bar, 10 nm scale bar for inset). C: TEM micrograph of negative stained released vesicles from neutral SMALPs (500 nm scale bar). D: Cryo-TEM micrograph of vesicle formed with neutral lipid SMALPs. A disturbance of the lipid bilayer is highlighted with a red circle (100 nm scale bar, 10 nm scale bar for inset). E: An example of a round vesicle observed during the cryo-TEM imaging of the sample in D (scale bar 20 nm). F: An example of aggregated vesicles observed during the cryo-TEM imaging of the sample in D (scale bar 100 nm).

To delve further into the structural characteristics of the vesicles generated through this SMALP assembly method and the electroformation method, cryo-TEM analysis was conducted. The results are presented in Figure 5B and D. In the case of electroformed vesicles, we observed relatively spherical structures characterised by an uninterrupted lipid bilayer with an approximate thickness of 5 nm. This bilayer thickness aligns with expectations derived from established literature sources.^38,39^ Conversely, vesicles formed from lipid SMALPs exhibited a more varied and less uniform morphology, occasionally featuring anomalous structures, as observed in Figure 5D-F. While a discernible lipid bilayer was still evident, it appeared less uniform and more disrupted in comparison to the electroformed GUVs. This structural difference may be attributed to the potential presence of residual polymer within the bilayer of the vesicle. Previous studies have demonstrated that even small quantities of polymer can lead to pore formation in lipid bilayers. These pores may subsequently compromise the stability of the formed vesicles,^40,41^ potentially contributing to the observed irregularities and deformations during the cryo-TEM sample preparation processes and explain the poor GUV release efficiency at high SMALP concentrations.

In order to further elucidate the characteristics of the vesicle membrane, a permeability study was conducted by introducing a fluorescein solution to the outer buffer around the vesicles and subsequently monitoring the extent of leakage (fluorescence intensity inside the GUVs) at the moment of introduction of the fluorophore and after 10 min. This investigation revealed that for all lipid compositions, fluorescein leaked into the interior of the vesicles, implying the existence of pores within the lipid bilayer of a dimension exceeding that of fluorescein molecules (Figure 6B).

**Figure 6:**
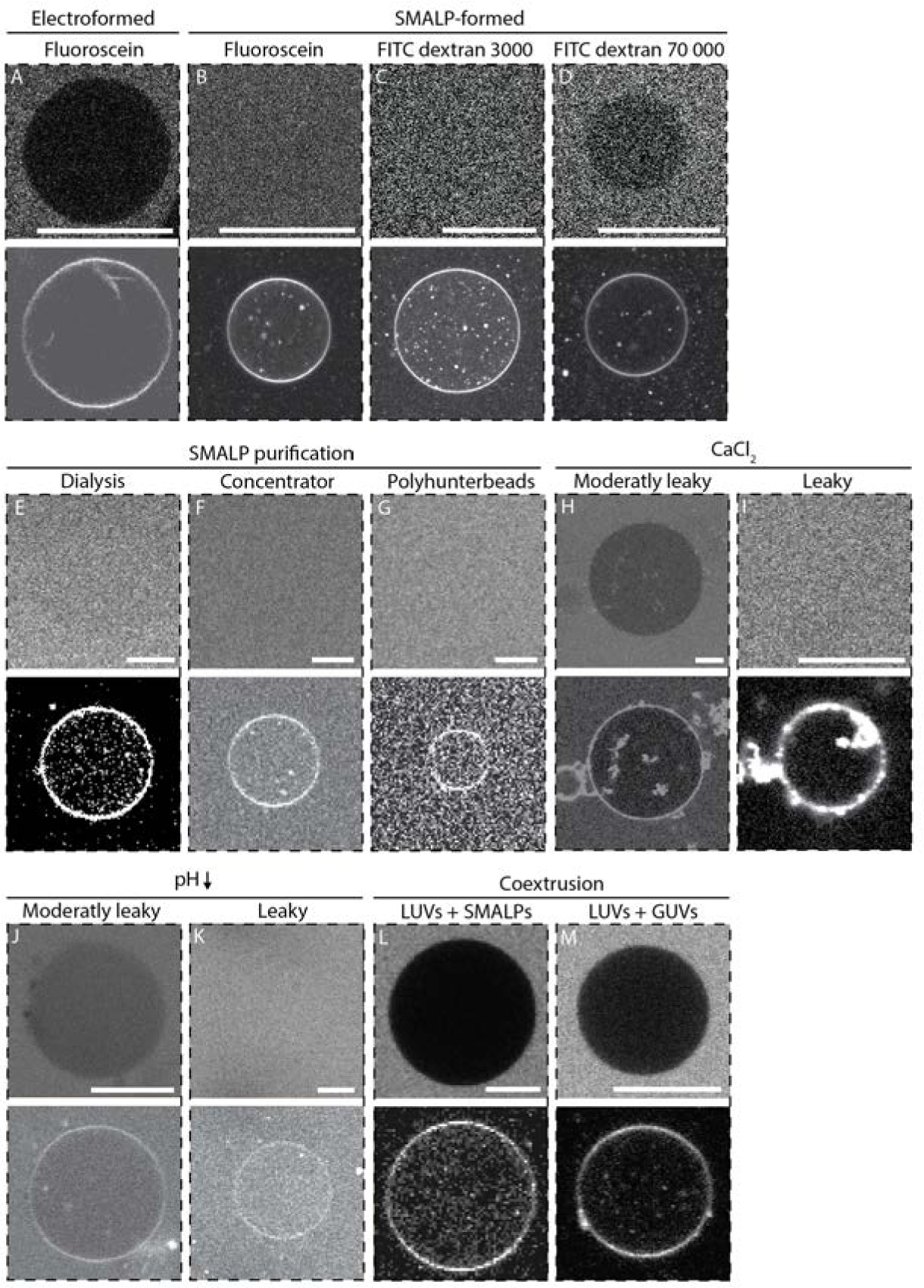
Fluorecence micrographs of leakage assay A-D and different routes for potential leakage mitigation E-M (top fluorescein channel, bottom rhodamine channel). A: Fluorescein added to electroformed GUVs. B: Fluorescein added to vesicles formed with neutral SMALPs (3.6 w/w% Krytox/1.4 w/w% RAN). C: FITC-dextran (MW 3000) added to vesicles formed with neutral SMALPs (3.6 w/w% Krytox/1.4 w/w% RAN). D: FITC-dextran (MW 70 000) added to vesicles formed with neutral SMALPs (3.6 w/w% Krytox/1.4 w/w% RAN). E-G: Fluorescein leakage assay of vesicle produced with a cleaned SMALP solution by dialysis (E), using a concentrator (F) and by polyhunterbeads (G). H-K: Fluorescein leakage assay of vesicle produced with SMALPs after addition of CaCl_2_ (H and I) or HCl (J and K) For both cases non-leaky and leaky vesicles were observed. L and M: vesicles produced with: a lipid solution formed by coextruding LUVs with SMALPs (L) and a lipid solutions formed by coextruding LUVs with vesicles formed from SMALPs (M). A-D: 20 µm scale bar, E-M: 10 µm scale bar.

Similar results were obtained for FITC-dextran with a molecular weight of 3000, but for larger fluorescent molecules, such as FITC-dextran with a molecular weight of 70 000, the rate of diffusion was decreased, as evidenced in Figure 6C-D. This outcome corroborates the aforementioned observation of a disturbed vesicle membrane characterised by discernible pores of variable sizes, a phenomenon also documented during cryo-TEM analysis.

In order to validate our hypothesis regarding the persistence of copolymer on the vesicle membrane, vesicles were generated using neutral FSMALPs without the incorporation of rhodamine lipids. Consequently, the sole source of fluorescence in these vesicles arises from the modified FSMA. The resulting image, displayed in Figure S10, distinctly exhibits individual vesicles, thereby substantiating the continued presence of FSMA on the vesicle membrane. This observation potentially provides an explanation for the observed permeability issues,^40,41^ underlining the significance of addressing and mitigating the residual polymer content in the vesicle formation process.

### Reducing vesicle leakiness

To alleviate the issue of vesicle permeability, several strategies were implemented. The initial approach centred on the removal of free SMA from the SMALP solution. This was guided by the hypothesis that trace amounts of unbound SMA within the vesicles could potentially interact with the lipid bilayer, thereby facilitating pore formation at low concentrations. Three distinct methodologies were employed to achieve this purification: dialysis, using a centrifugal concentrator, and the utilisation of polyhunter beads (a product developed by Cube Biotech for removal of free polymer solution). Following each purification process, vesicle permeability was assessed via a fluorescein leakage assay (Figure 6E-G).

In all three instances, fluorescein leaked instantly into the vesicles, suggesting that the exclusive removal of free SMA did not yield the desired outcome in terms of mitigating vesicle leakage. This observation underscores that the polymer encompassing the SMALPs themselves is the primary source, persisting on the vesicle surface.

In subsequent experiments, attempts were made to eliminate the polymer presence on the surfaces of GUVs through two distinct approaches: Introduction of divalent cations (Ca^2+^) and reduction of pH through HCl Addition. Ca^2+^ has been reported to induce polymer aggregation in literature.^27,42,43^ Visible aggregation of lipid structures occurred after Ca^2+^ addition. This led to the disruption of some GUVs; nevertheless, a subset of GUVs remained intact. Alternatively, SMA can be aggregated by a reduction of the pH as previously reported.^32,44,45^ We reduced the pH by introducing HCl into the system, thereby aggregating the polymer. As with the Ca^2+^ addition, changes in buffer conditions were implemented following vesicle production. Similar to the Ca^2+^ experiment, visible lipid aggregation was observed post-condition modification, resulting in the destruction of some GUVs. Notably, after these interventions, some of the GUVs exhibited reduced leakiness, as evidenced by a discernible dark sphere in the fluorescein channel. However, this improvement proved to be transient, as leakage was still observable within less than 10 minutes, signifying that the vesicles retained their leakiness characteristics (see Figure 6H-K). Furthermore, a subset of the vesicles displayed rapid leakage as observed prior to the introduced modifications.

For the final set of experiments, endeavours were undertaken to first convert SMALPs into LUVs, from which any residual SMA was washed away via centrifugation. Two distinct techniques were employed. In the first approach, lipid SMALPs were subjected to a coextrusion process alongside LUVs. Subsequently, the resulting LUVs underwent a rigorous washing process aimed at removing any remaining free SMA. It was shown in earlier research that proteins could be incorporated in LUVs via this route.^46^ The second approach involved the generation of leaky vesicles from SMALPs. These vesicles were subsequently collected, released, and subjected to a coextrusion process alongside another LUV solution, devoid of SMA. Post-coextrusion, the resultant LUVs were washed by centrifugation to eliminate any lingering SMA. Remarkably, both methodologies yielded non-leaky vesicles, as evidenced in Figure 6L-M. As such, these routes provide evidence of the importance of effectively removing free SMA from the outer GUV buffer to obtain suitably compartmentalising vesicles. An important remark regarding this approach, is that the transfer efficiency of proteins stabilised in SMALPs into the LUVs is relatively low,^47^ and therefore a more direct strategy for the formation of GUVs directly from SMALPs would prove useful in the future.

## Conclusion

In this study we successfully demonstrate that SMALPs can be assembled at a droplet interface to form GUVs, similarly to the method developed by Weiss *et al.*^19^ which utilises LUVs. The assembly process results from the interaction, at the droplet interface, between the SMALP-forming polymer and the carboxylic groups present in the Krytox surfactant. This interaction was confirmed via plate assays and the encapsulation of free polymer within droplets. To facilitate monitoring of these interactions, a novel SMA variant, FSMA, was developed, incorporating fluorescein for tracking the distribution of the SMA polymer within surfactant-stabilised droplets.

Furthermore, we generated vesicles from SMALPs with a wide array of lipid compositions, pH values, SMA concentrations and surfactant mixtures. Our experiments revealed that, in contrast to methods utilising LUVs,^37^ our method enabled the formation of vesicles from various lipid compositions, independent of lipid charge. We further report that the release efficiency of vesicles from their stabilising droplet environment diminished at higher pH levels, and we identify an optimal Krytox and SMA concentration for vesicle assembly.

Additionally, vesicle integrity was comprehensively assessed using (cryo-)TEM imaging and a fluorescein leakage assay. TEM images unveiled vesicle-like structures, distinct from a mere aggregation of SMALPs. Nevertheless, closer examination by cryo-TEM revealed interruptions in the lipid bilayer, which were attributed to the presence of intercalated SMA polymer molecules. Importantly, fluorescein leakage assays signalled the presence of pores large enough to permit the traffic of small molecules like fluorescein, but small enough to prevent passage of larger molecules (such as FITC-dextran, 70 000 g/mol). Employing the newly developed FSMA, we observed the persistence of FSMA on the vesicle surface, providing an explanation for the potential role of SMA pores (the formation of which has previously been established^40^) on vesicle leakiness.

Several strategies were explored to mitigate vesicle leakiness by removing free SMA polymer from the SMALP samples, such as dialysis, the use of a concentrator and treatment with polyhunter beads. Regrettably, these efforts did not yield substantial improvements. Notably, the addition of Ca^2+^ and pH reduction led to SMA polymer aggregation, resulting in less permeable vesicles, albeit leakage was still observed. A strategy that proved effective in reducing vesicle leakage consisted in the transfer of SMALPs into LUVs through coextrusion with a lipid solution, followed by the utilisation of these LUVs to form the final GUVs. This problem warrants further exploration and investigation, since direct GUV formation from SMALPs could provide a route to better preserve the functionality of inserted membrane proteins by circumventing any detergent mediated steps. A potential solution could come from the use of alternative nanodisc-forming polymers,^48–51^ which may be more amenable to removal after GUV formation.

## Experimental

### Materials

Ammonium molybdate(VI) tetrahydrate solution, Ascorbic acid, H_2_SO_4_, cholesterol, dimethylsulfoxide (DMSO), fluorescein sodium salt, glucose, NaCl, PEG 2000, perfluoro-1-octanol, sucrose, tris(hydroxymethyl)aminomethane (Tris), 1,2-dioleoyl-sn-glycero-3-phosphocholine (DOPC), 1-palmitoyl-2-oleoyl-glycero-3-phosphocholine (POPC), 1,2-dioleoyl-3-trimethylammoniumpropane (chloride salt) (DOTAP), 1,2-Dioleoyl-sn-glycero-3-phopsho-(1’-rac-glycerol (sodium salt)) (DOPG), Egg phosphatidylcholine (EggPC), Egg phosphatidylglycerol (EggPG), 1,2-dioleoyl-sn-glycero-3-phosphoethanolamine-N-(lissamine rhodamine B sulfonyl) (ammonium salt) (18:1 Liss Rhod PE) were purchased from Sigma–Aldrich (Germany). Fluorescein dextran 3000 and 70 000 from ThemoFisher (UK). RAN 008 was acquired from RAN Biotechnologies (US), Krytox 157 FSH from Costenoble (Germany). The fluorintated oil HFE 7500 from Fluorochem (UK). Ultrapure water was produced with the Milli-Q SynErgy UV system from Merck Millipore. Erythrocytes were obtained from Red Cross Flanders.

## Methods

FSMA was prepared reacting SMA2.3 with putrescin-FITC (see Supplementary material for a detailed protocol).

### FSMALP/SMALP formation

Lipid solutions were prepared in chloroform for four different lipid compositions (neutral, negative-1, negative-2, positive) with the following compositions (% in mol percentage). Neutral: 99.5% DOPC, 0.5% liss rhod PE, negative-1: 34.75% DOPC, 34.75% POPC, 15% DOPG, 15% Cholesterol, 0.5% Liss rhod PE, negative-2: 80% EggPC, 19.5% EggPG, 0.5% Liss rhod PE, positive: 45% DOPC, 45% POPC, 9.5% DOTAP, 0.5% Liss rhod PE. Afterwards, lipids were dried by flushing with N_2_. Subsequently, the lipid film was further desiccated for an additional 2 h to eliminate any residual chloroform. The lipid films were rehydrated with a SMALP buffer solution (30 mM Tris, 300 mM NaCl and pH 7.5). Lipids were vortexed for 30 s and subjected to bath sonication for 30 min. Polymer solution, either SMA or FSMA, was added in a ratio of 2 w/v% polymer solution for 42 mg/ml membrane solution if not stated otherwise. The polymer-lipid mixture was allowed to incubate for a minimum of 2 h at room temperature. Subsequently, FSMALPS/SMALPs were isolated from any remaining debris by centrifuging at 100 000 g for 45 min.

### Vesicle and droplet imaging

Droplet samples were imaged on a Nikon Ti2 fluorescence microscope (Nikon, Japan) by transferring 8 µl of droplet solution into a cell counting chamber from Kisker Biotech GmbH (Germany). Vesicle solutions were imaged by transferring 50 µl of solution into 50 µl of SMALP buffer solution in one of the wells of a 16 well uncoated polymeric Ibidi slides (Ibidi GmbH, Germany).

### FTIR

SMA and FSMA were first precipitated with HCL and then thoroughly washed three times with DI water. Next the polymers were frozen in liquid nitrogen and lyophillised into a dry powder. The powder was transferred to the ATR-FTIR and spectra were measured in transmittance from 400-4000 cm*^−^*^1^.

### DLS

DLS measurements were conducted using the Zetasizer Nano ZSP instrument (Malvern Panalytical, UK). The experiments were carried out in a quartz cuvette (ZEN 2112) at an angle of 173° (25°C) in triplicate. The following material properties were employed: SMALP buffer (refractive index: 1.330, dynamic viscosity: 0.887 cP), lipids (refractive index: 1.450), silica particles (refractive index: 1.540).

### TEM

A 300 mesh copper TEM grid was glow discharged for a duration of 15 s. Then 3.5 µl of 100x diluted sample was deposited on the grid and incubated for 5 min. Subsequently, for the negative staining, 30 µl of 1 v/v% uranyl acetate was incubated on the grid for 1 min. Finally, samples were blotted and imaged.

### Plunge-freezing and cryo-TEM

3.5 µL of a sample of GUVs was deposited on a lacey grid, glow-discharged in a Leica ACE600 (Leica, Vienna, AT) coating unit, and held by a forceps in a humidity controlled chamber of a Leica GP2 plunge-freezer. Then, after back-blotting for 2 s, the grids were vitrified by plunging in liquid ethane close to its freezing point, and stored under liquid nitrogen. Samples were observed and imaged in a JEOL F200 (JEOL, Tokyo, JP) transmission electron microscope, equipped with a Gatan Continuum energy filter and K3 camera, using zero loss filtering with a slid width of 20 eV. Images were taken with pixel size of 0.53 nm and a maximum exposure dose of less than 60 electrons per Å^2^.

### Lipid solubilisation

FSMALPs/SMALPs were formed in accordance with the previously described protocol utilising varying amounts of SMA or FSMA (0.06, 0.12, 0.18, 0.24, 0.37, 0.5, 1, 1.5 w/v% in a final of 6 mM neutral lipid solution). The lipid concentration within samples was determined through the analysis of the total phosphorus content (experiment performed in triplicate).

A set of five standard tubes were prepared with 0, 0.0325, 0.065, 0.114, 0.163, 0.228 µmol phosphorus standard, respectively, in a 2 ml glass vial. Additionally, sample tubes were prepared with 50 µl of sample. To both sets 11 µl of 8.9 M H_2_SO_4_ was introduced. All tubes were heated in an aluminium heating block at 200-215 °C for 25 min. Following heating, the tubes were removed and allowed to cool for 5 min after which 37.5 µl of H_2_O_2_ was added. Tubes were then subjected to an additional 30 min heating step. Afterward, the tubes were allowed to return to room temperature. Next, to all tubes 975 µl of ultrapure water and 125 µl of a 2.5 m/v% ammonium molybdate(VI) tetrahydrate solution was added. Tubes were vortexed 5 times. Following the ammonium molybdate addition, 125 µl of a 10 m/v% ascorbic acid solution was transferred to the tubes, which were vortexed another 5 times. The tubes were capped with aluminium foil and heated at 100 °C for 7 min. Subsequently, samples were cooled to ambient temperature and their absorbance was measured at 820 nm. A calibration curve was constructed using the phosphate standard, and the phosphate concentration in each sample was calculated.

### FSMALPs/SMALPs preparation from red blood cells

Production of FSMALPs/SMALPs derived from red blood cells (RBCs) was based on the protocol outlined by Desrames *et al.*^25^ The procedure is summarised as follows: 500 µl of pelleted RBC solution (RBCs from Red Cross Flanders) were washed 3 times with 1 ml of DPBS solution through centrifugation at 1000 g for 5 min. To 200 µl of RBCs 8 ml of low ionic strength buffer (LIS: 5 mM sodium phosphate, 0.1 mM EDTA, pH 8) was added and incubated for 15 min at 4 °C. The sample was then washed until a white pellet was obtained by centrifuging for 15 min at 27 000 g at 4 °C to replace the LIS solution. Subsequently, 8 ml of very low ionic strength buffer (VLIS: 0.3 mM sodium phosphate, 0.1 mM EDTA, pH 8) was added and sample was centrifuged for 15 min at 27 000 g at 4 °C. The supernatant was discarded and the sample was resuspended in 2 ml VLIS buffer, which was then incubated for 30 min at 37 °C for spectrin removal. VLIS buffer was removed and sample was resuspended in 200 µl of SMALP buffer to which 100 µl or 50 µl of 6 w/v% FSMA/SMA solution was added. The solution was rotated on a wheel for 2 h at 37 °C and further rotated overnight on the wheel in the fridge. At last the sample was centrifuged for 45 min at 100 000 g to remove all debris and FSMALPS/SMALPs were transferred to another centrifuge tube and further stored in the freezer for subsequent use.

Protein incorporation was assessed using western blot analysis specific for band 3 proteins, an abundant membrane protein of the RBC. For performing this experiment samples were run on a 7.5% gel, using as control a western blot for the cytosolic protein carbonic anhydrase 1 (CA-I), in which samples were run on a 12% gel.

### Krytox-FSMA interaction

**UV-VIS measurements**: The interaction between FSMA/SMA with Krytox was investigated by depositing 100 µl of a 0.08 w/v% FSMA/SMA in SMALP buffer solution on top of different Krytox solutions (10, 5, 1, 0.1, 0 w/w% Krytox). Solutions were left overnight and 50 µl of the oil phase was transferred to a 384 well plate (UV-STAR for SMA case and black well for FSMA). Absorbance (for SMA) was measured at 270 nm and fluorescence (for FSMA) at 470/550 (ex/em) nm with a spectrophotometer (SpectraMax ID3), conducting measurements in triplicate.

**FITC/FSMA/FSMALP encapsulation**: Fluorescein isothiocyanate was initially solubilised at 2 mg/ml in DMSO after which it was diluted 100x in SMALP buffer solutions. 50 µl of this solution was transferred on top of 100 µl of the three different surfactant solutions: 1 w/w% RAN, 3.6 w/w% Krytox/1.4 w/w% RAN and 5 w/w% Krytox/1.4 w/w% RAN respectively. Droplets were vortexed for 15s and imaged. FSMA and FSMALPs were similarly encapsulated, FSMA at a concentration of 0.08 w/v% and FSMALPs at lipid a concentration of 1 mM.

### GUV production

SMALPs were generated from the four distinct lipid mixtures and were encapsulated in droplets stabilised by two different surfactant formulations. Formulation 1: 3.6 w/w% Krytox/1.4 w/w% RAN and formulation 2: 5 w/w% Krytox/1.4 w/w% RAN. 50 µl of SMALP solution (*±* 1 mM) was deposited on top of 100 µl of surfactant solution. The resulting mixture was vortexed for 30 s and the formed droplets were incubated overnight in the fridge allowing the SMALPs to interact with the Krytox polymer. Vesicles were released by adding 50 µl of extra buffer solution and 50 µl of 20 v/v% PFO solution. Vesicles were then transferred to an Ibidi imaging chambers.

Additionally, the effects of pH, Krytox concentration and SMA concentration for neutral SMALPs were investigated by slightly altering the protocol above. For the effect of the pH on neutral SMALPs was investigated by using a slightly adjusted SMALP buffer (300 mM NaCl and 30mM Tris) at pH 8 and 9. And the effect of Krytox concentration by using additional surfactant mixtures 10 w/w% Krytox, 7 w/w% Krytox/1.4 w/w% RAN, 1 w/w% Krytox/1.4 w/w% RAN and 1 w/w% RAN besides the standard 5 w/w% Krytox/1.4 w/w% RAN and 3.6 w/w% Krytox/1.4 w/w% RAN. For investigating the effect of polymer concentration different concentration of SMA (0.12, 0.24, 0.48, 0.96 w/v% in a 6 mM neutral lipid solution) were used to make SMALPs, these SMALPs were then encapsulated in droplets and released in later experiments.

The integrity of the produced vesicles was tested by introducing 10 µl of a 0.7 µM fluorescein solution to 100 µl of vesicle solution. The samples were imaged over time just after addition and after 10 min. Additionally, experiments were performed with FITC-dextran with a molecular weight of 3000 and 70 000.

### Removal of free SMA

The removal of free SMA polymer from SMALP solution was performed with three different techniques: dialysis, utilising a concentrator and flowing the SMALP solution over polyhunterbeads (Cube Biotech GmBH, Germany). In all cases SMALP solution was produced as described earlier.

**Dialysis**: SMALP solution was dialysed against a 50 KDa dialyis membrane against a 100x volume excess (of SMALP buffer) for 1 h, followed by an additional 3 h dialysis against a 100x volume excess and an overnight dialysis.

**Concentrator**: A 100 kDA concentrator was first washed with a 5 w/v% PEG 2000 solution. After which the 2x diluted sample was added and upconcentrated to its original concentration. This procedure was performed three times.

**Polyhunterbeads**: These special beads are developed by Cube Biotech (Germany) for free polymer removal. 2 ml of the polyhunter solutions was transferred to a gravity flow column. Beads were washed 3 times with distilled water and two times with buffer solution. 0.7 ml of a 6 mM lipid solution was added and eluted with the SMALP buffer solution.

SMA polymer solution is known to aggregate at high CaCl_2_ and low pH conditions.

**CaCl_2_ precipitation**: 50 µl of vesicle solution was deposited in a well prefilled with 50 µl of SMALP buffer. To this solution 50 µl of a 150 mM NaCl, 100 mM CaCl_2_ and 30 mM Tris (pH: 7.5) solution was added.

**Reduction of pH**: Here instead of adding the CaCl_2_ solution 1 µl of a 20 mM HCl solution was transferred to the vesicle solution. These conditions were utilised in the following procedures as a route for removing excess polymer.

For both protocols samples were incubated for 5 min before performing the fluorescein leakage assay.

**Coextrusion of SMALPs with LUVs**: 990 µl of a 2 mM lipid solution was coextruded with 10 µl of a 2 mM SMALP solution through a 100 nm polycarbonate membrane with a mini extruder (Avanti Polar Lipids, USA). Resulting LUVs were washed 2 times by centrifugation (100 000g for 30 min).

**Coextrusion of vesicles with LUVs**: Vesicles were produced as described earlier, 250 µl of this vesicle solution was combined with 250 µl of a 2 mM LUV solution and extruded through a 100 nm polycarbonate membrane. The resulting LUVs were washed by centrifugation as described in the previous section.

Lipid solutions were produced through the same protocol as SMALPs formation until the step just before SMA polymer addition.

### Electroformation

A total of 10 µl of a 1.5 mg/ml lipid chloroform (DOPC) solution was deposited onto the conductive side of an indium tin oxide (ITO) coated slide. This slide was left to desiccate overnight, ensuring removal of residual chloroform. Next a rubber ring was affixed around the dried lipid spot using a wax seal. The lipids were then hydrated by gently adding 250 µl of a 300 mM sucrose solution. A second ITO coated slide was placed on top. These two slide were transferred to the ‘Vesicle prep Pro’ device (Nanion, Germany). 3 V was applied to the bilayer at a frequency of 5 Hz for a duration of 160 min. During the electroformation the temperature was gradually increased from room temperature until 36 °C over 30 min. The voltage was incrementally increased, first from 0.1 to 0.5 V over 30 min and then to 3 V over 15 min. In the end the voltage was gradually decreased to 0 V over 5 min. GUVs were afterwards collected with a wide orifice pipette tip and stored in the fridge.

## Acknowledgement

This research has received funding from the Research Foundation Flanders with grant No 1S43521N and No G074321N. Additionally, this work received funding from the European Union’s Horizon Europe research and EIC grant agreement No 101046894 (SynEry) and No 101130715 (ArTCell). Views and opinions expressed are however those of the author(s) only and do not necessarily reflect those of the European Union or the granting authority European Union’s Horizon Europe research and innovation program. Neither the European Union nor the granting authority can be held responsible for them.

## Notes

### Competing Interest Statement

The authors have declared no competing interest.

