## Supplementary material for "Lipid vesicle formation by encapsulation of SMALPs in surfactant-stabilised droplets"

### Supporting Information Available

#### FSMA preparation

In a round bottom flask, 75 mg (0.19 µmol, 1 eq) of Fluorescein diacetate 5(6)-isothiocyanate, 33 mg (0.21 µmol, 1.1 eq) of N-Boc-putrescine and 79 µL of triethylamine (0.57 µmol, 3 eq) were dissolved in DMF (1 mL). The whole was stirred at room temperature until the reaction completion, monitored by TLC. The crude was reacetylated with acetic anhydride and pyridine and purified by HPLC yielding 20 mg of solid.

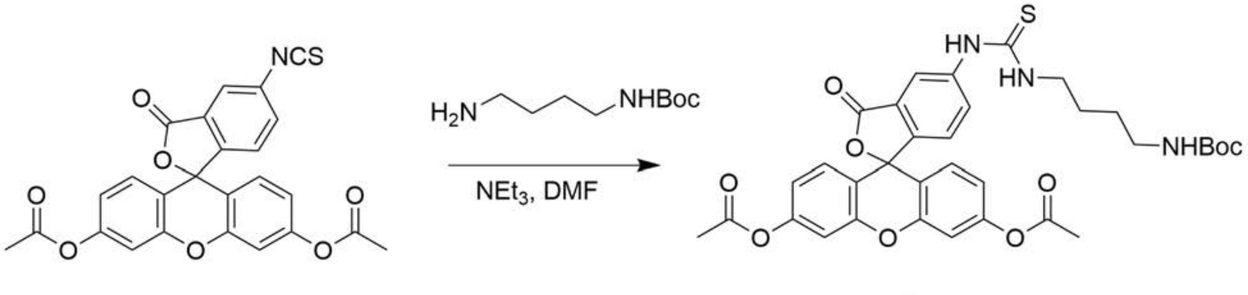


Figure S1: Synthesis of 5-(3-(4-((tert-butoxycarbonyl)amino)butyl)thioureido)-Fluorescein diacetate, **2**.

In a round bottom flask, SMA2.3:1 (1 eq), 5-(3-(4-((tert-butoxycarbonyl)amino) butyl)thioureido)-Fluorescein diacetate (2 eq), HATU (2.4 eq) and DIPEA (1.9 eq) were dissolved in DMA and stirred overnight. After this time, the whole was acidified using HCl to pH 2 resulting in FSMA precipitation. FSMA was washed, purified and solubilized in a 6 w/v% working solution in 50 mM HEPES 300 mM NaCl pH 8, as previously described.[52]

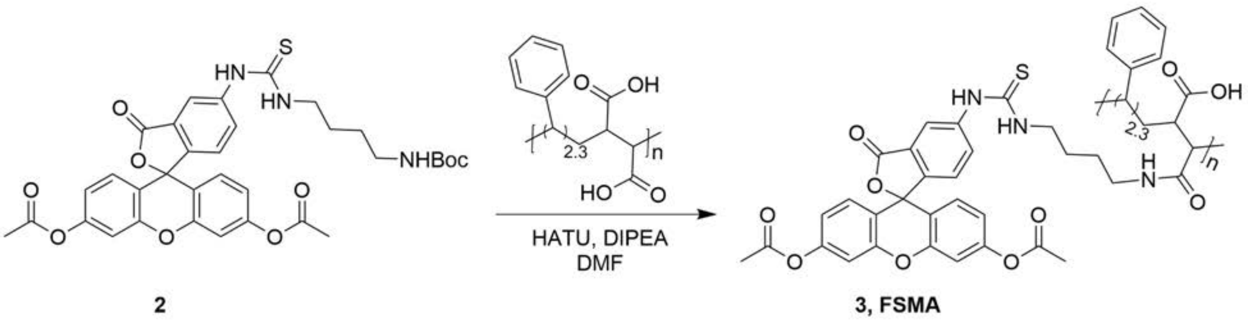


Figure S2: Synthesis of FSMA, **3**.

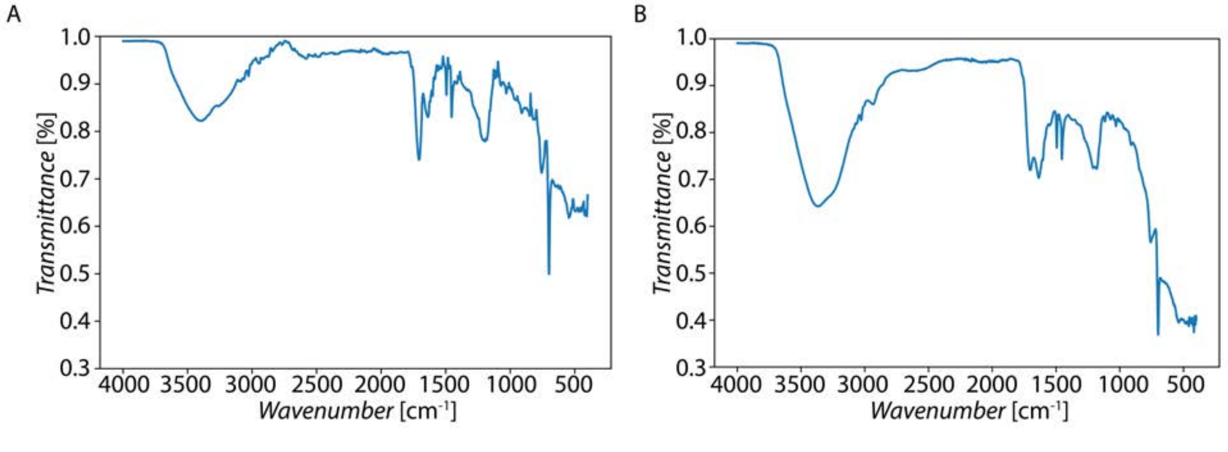


Figure S3: FTIR spectra of A: SMA and B: FSMA.

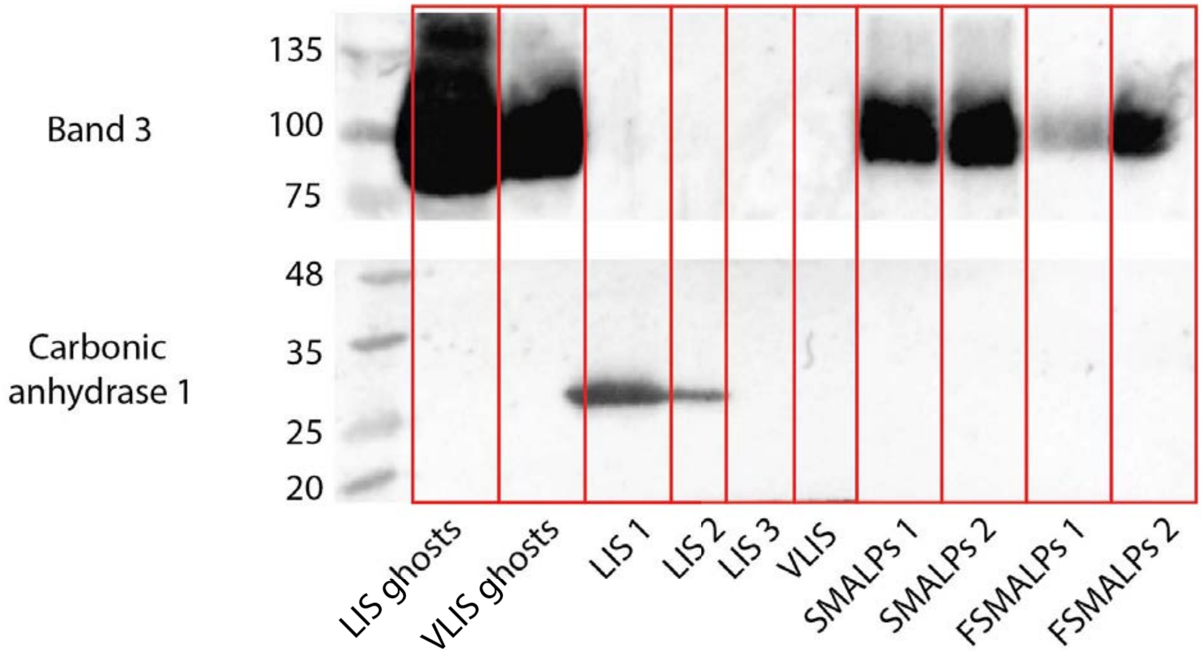


Figure S4: Western blot for band 3 and carbonic anhydrase 1. LIS ghosts: RBC ghosts obtained after LIS washes. VLIS ghosts: RBC ghosts obtained after VLIS washes. LIS/VLIS supernatant obtained after the respective VLIS/LIS washes. SMALPs and FSMALPs are the final nanodiscs, 1 with 50 µl of 6 w/v% polymer solution added, 2 with 100 µl of 6 w/v% polymer solution added.

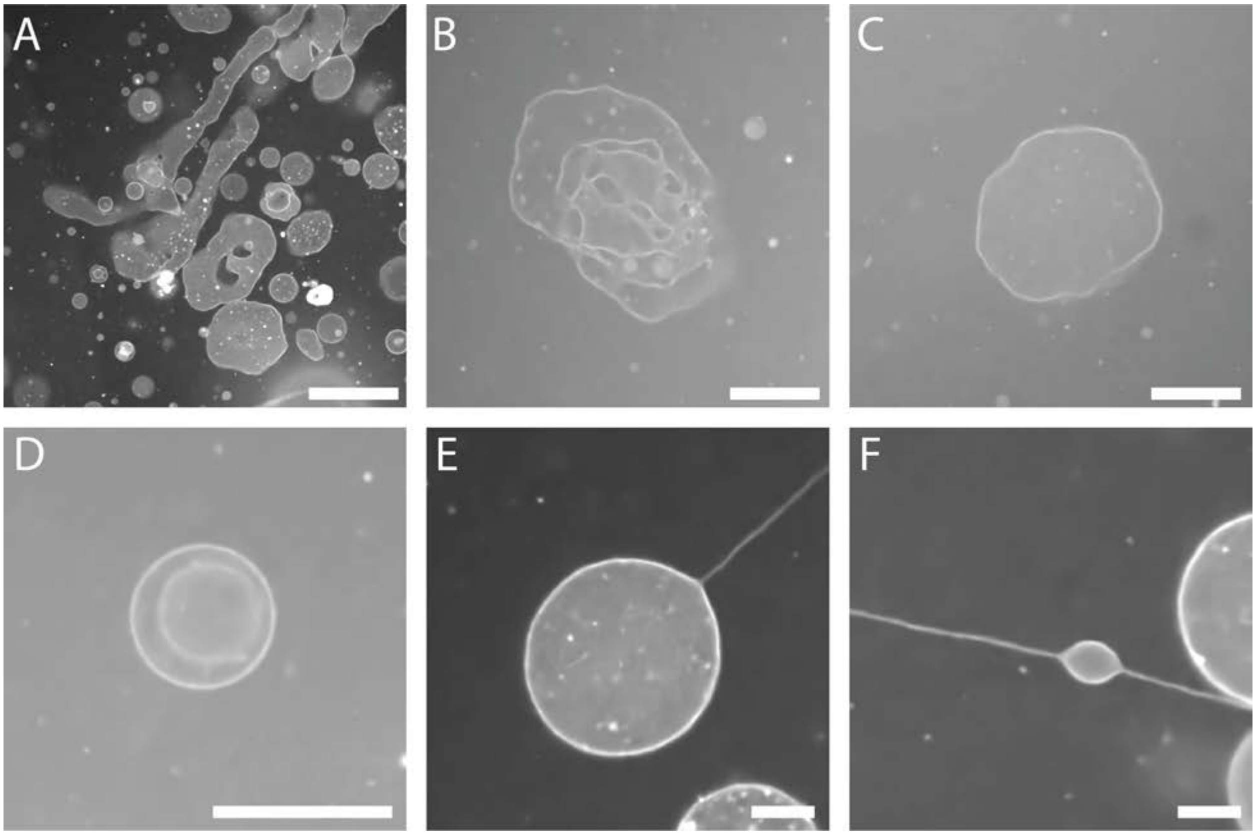


Figure S5: Collection of the special effects of SMA on vesicle formation. A: Long elongated vesicles sometimes stuck to imaging chamber surface (50 µm scale bar). B: Collapsed vesicles (25 µm scale bar). C: Partially deflated vesicle (25 µm scale bar). D: Vesicle in vesicle (25µm scale bar). E and F: Formation of thin tubulation (25 µm scale bar).

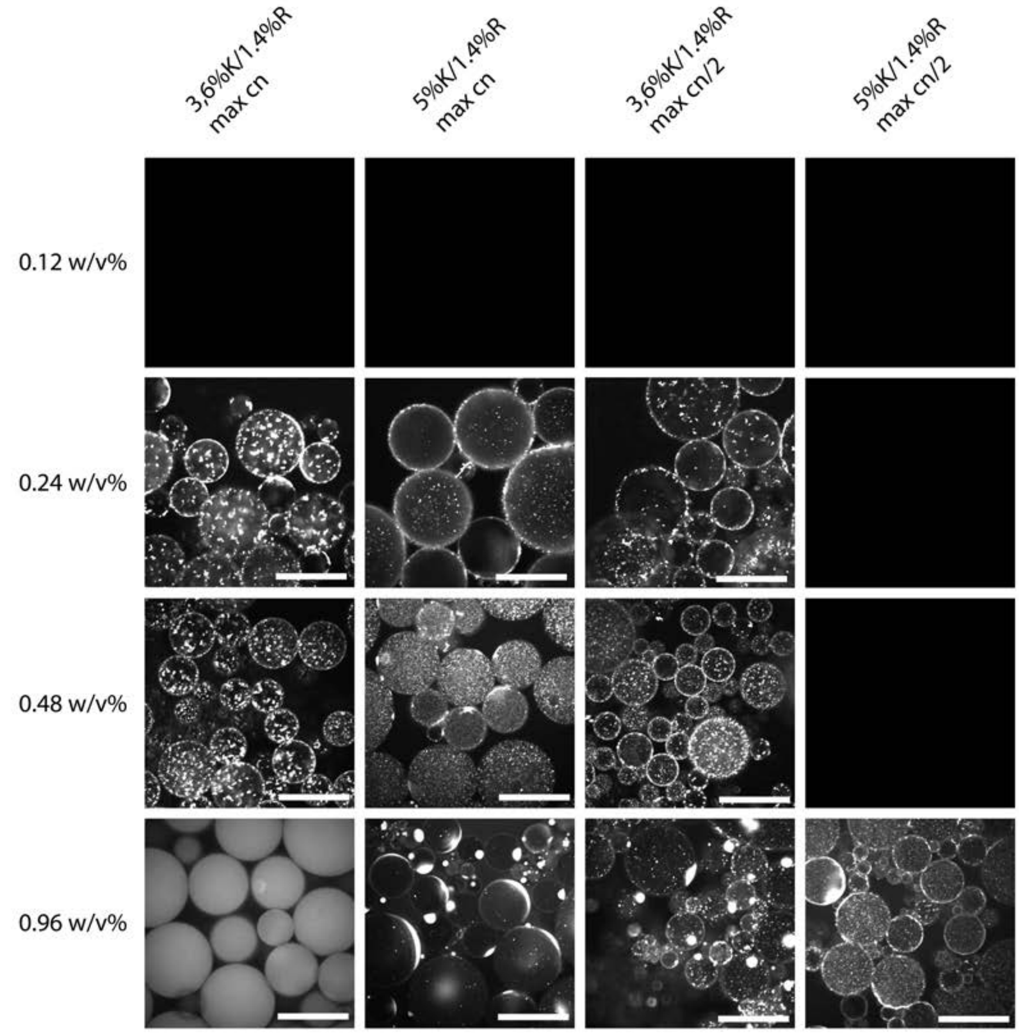


Figure S6: Fluorescent micrographs of droplets formed with the neutral lipid SMALPs formed with different amounts of SMA and stabilised by different surfactant mixtures at two different SMALP concentrations (max: is the concentration of undiluted SMALPs, max/2 is 1/2 diluted). Black panels indicate unstable droplets which could not be imaged (100 µm scale bar).

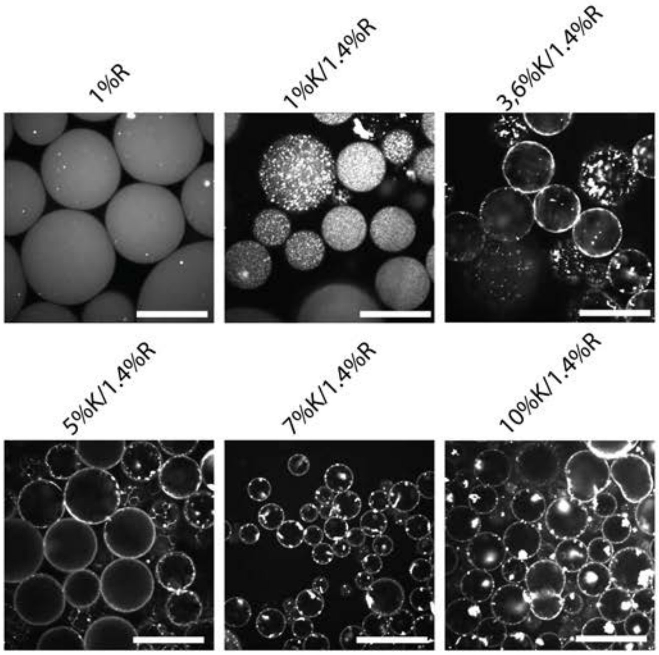


Figure S7: Fluorescent micrographs of droplets encapsulating neutral lipid nanodiscs at 1 mM for different surfactant mixtures.

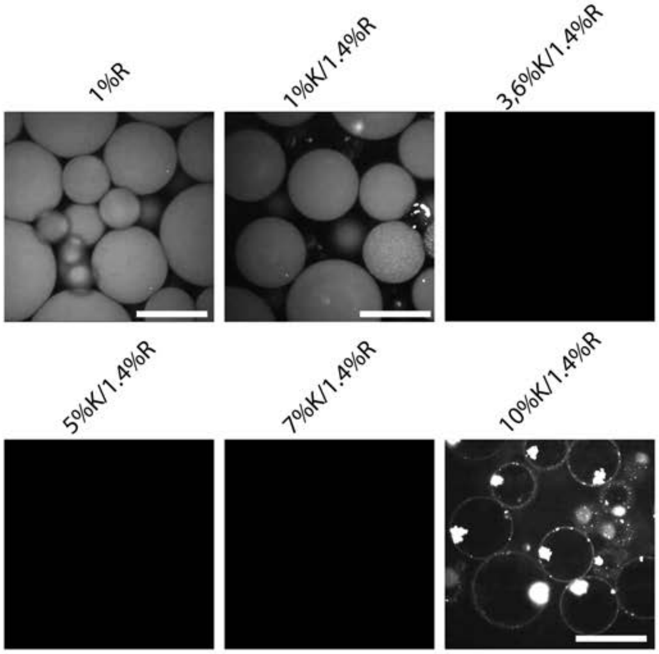


Figure S8: Fluorescence micrographs of droplets encapsulating neutral lipid SMALPs at 0.5 mM for different surfactant mixtures.

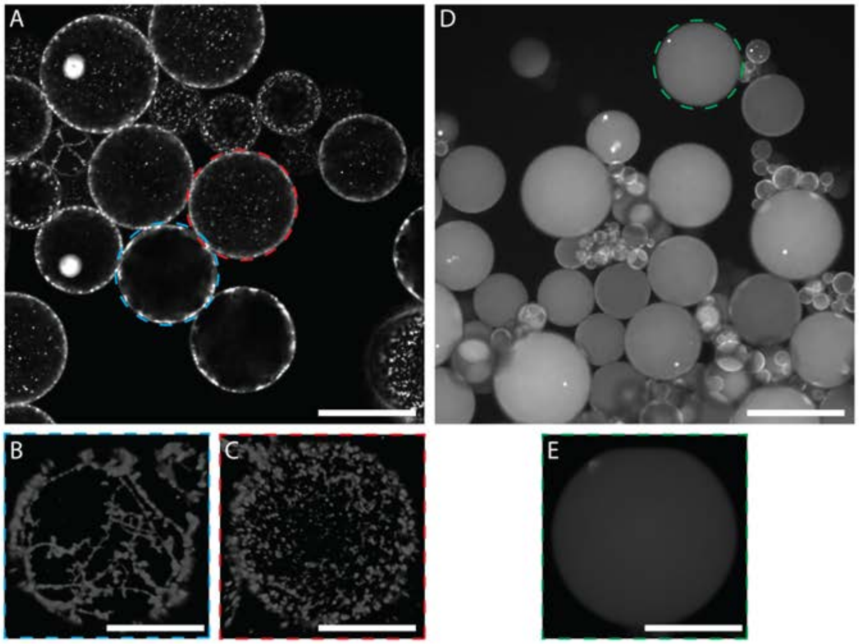


Figure S9: A: Confocal image of droplets stabilised by 5 w/w% Krytox and 1.4 w/w% RAN with neutral lipid SMALPs at pH 8. A 3D rendered Z-stack is shown in panel B and C (A: 100µm scale bar, B and C: 50 µm scale bar). D: Confocal image of droplets stabilised by 5 w/w% Krytox and 1.4 w/w% RAN with neutral SMALPs at pH 9. E: A 3D rendered Z-stack of a droplet in D (D: 100 µm scale bar, E: 50 µm scale bar).

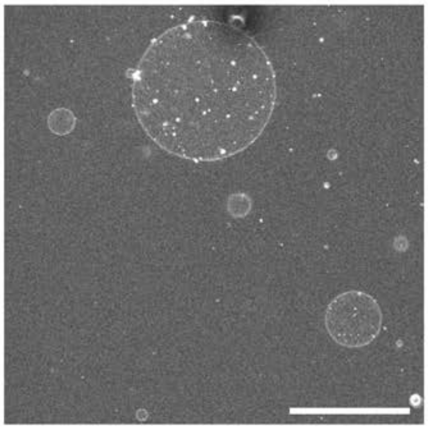

Figure S10: Vesicles formed with FSMALPs no fluorescent lipids present imaged in fluorescein channel (50 µm scale bar).
